# Production of P2D1 tailocins by *Dickeya dadantii* 3937: a temporal relationship between the stressor onset, gene expression, and the concentration of active particles

**DOI:** 10.1101/2024.11.21.624640

**Authors:** Marta Sobolewska, Dorota M. Krzyżanowska, Marcin Borowicz, Robert Czajkowski

## Abstract

Tailocins are bacteriocins resembling bacteriophage tails. Previously, we reported the production of P2D1 tailocins in the plant pathogen *Dickeya dadantii* 3937, the synthesis of which was upregulated upon treatment with mitomycin C. In the current study, we characterized the temporal relationship between the inducer onset, activation of expression of tailocin-related genes, the concentration of tailocin particles, and the titer of viable producer cells. Using a newly established RT-qPCR assay, we found that the expression of P2D1 structural genes peaks two hours after the addition of mitomycin C. Simultaneous measurement of the titer of tailocins showed that the maximum concentration of these particles in the culture supernatant was reached 6 hours post induction and remained stable till the termination of the experiment (18 hours). Progressing accumulation of tailocins between the second and the sixth-hour post mitomycin C treatment correlated with a sharp drop in the count of viable cells (ca. five orders of magnitude). This effect was related to the mechanism by which tailocins are released from the cells, which involves cell lysis. Our study is the first detailed analysis of the timing of events during P2D1 tailocin synthesis in *Dickeya* spp.

## Introduction

Approximately 70-80% of sequenced bacterial genomes contain genes of viral origin, allowing the holders to utilize some of these components for diverse functions independently from phage-bacteria interactions ^1,2^. Among these various genetic elements, some can encode for producing phage tail-like particles known as tailocins ^3^. Tailocins are nanomolecular structures resembling syringes that share evolutionary and morphological features with bacteriophage tails, type VI secretion systems, and extracellular contractile injection systems ^4^. These phage tail-like particles are used as defense mechanisms or offensive tools by bacteria ^5^. They are released in response to environmental stresses or competitive pressure, acting as precision weapons, targeting and killing kin bacterial competitors ^6^. By eliminating those closely related strains, the producers improve their chances of survival and resource acquisition in various ecological niches ^7,8^.

Tailocin production has been described both in Gram-negative and Gram-positive bacteria. The producing strains encompass a range of human, animal, and plant pathogens as well as saprophytic bacteria found in diverse environments, including rhizosphere, bulk soil, plant, water, insects, animals, and humans (reviewed in: ^9^). Likewise, we recently described a novel tailocin named dickeyocin P2D1 produced by the plant pathogenic bacterium *Dickeya dadantii* strain 3937. P2D1 tailocins resembled the tails of the bacteriophage *Peduovirus* P2. They exhibited bactericidal activity specifically against phylogenetically related bacterial strains without causing any adverse effects on eukaryotic cells in a *Caenorhabditis elegans* killing model ^10^.

Genetic cluster encoding P2D1 tailocin seems to be widely distributed in the genomes of *Dickeya* spp. In our recent screen of 74 complete and high-quality *Dickeya* spp. genomes deposited in the NCBI database, 52 genomes (70% of the genomes) contained tailocin clusters expressing high homology to the P2D1 cluster (>70% threshold for coverage and identity) ^11^. The fact that P2D1 tailocins are ubiquitously present in this group of bacteria suggests they likely confer a competitive advantage in kin interactions.

Production of tailocins is known to be triggered by the SOS response due to DNA damage ^12^. Still, the exact mechanism of induction and production of phage tail-like particles remains poorly understood for most analyzed strains. Under laboratory settings, the most commonly used inducers of phage tail-like particles are UV radiation and mitomycin C. UV radiation primarily induces reversible thymine dimers in DNA. This activation triggers the cleavage of the LexA repressor, leading to the derepression of SOS-regulated genes, including those involved in the production of tailocins ^13^. In contrast, mitomycin C is a DNA crosslinking agent that induces severe and irreversible DNA damage, activating the SOS response ^14^. Mitomycin C may cause a stronger and more prolonged SOS response compared to UV radiation and, therefore, probably elevated production of tailocins compared to UV radiation *in vitro* ^15^.

Among the current knowledge gaps are the environmental conditions under which the production of tailocins is triggered and how these conditions may contribute to the overall success of a particular strain in a given niche ^6^. Additionally, there is limited data on the timeline of events between the induction of tailocin production and the accumulation of active particles. Therefore, our study aimed to analyze the conditions and triggers of P2D1 production in *D. dadantii* strain 3937. Specifically, we explored the mitomycin C concentration-dependent and time-dependent production of P2D1 and analyzed the expression of P2D1 genes upon induction in real time using qRT-PCR. Likewise, we detailed other triggers of P2D1 induction and discussed how these compounds could modulate the concentration of P2D1 tailocins under environmental conditions.

## Materials and Methods

### Bacterial strains, chemical compounds, and growth media

*Dickeya dadantii* strain 3937 ^16^ and *Musicola paradisiaca* strain NCPPB 2511 ^17,18^ were grown at 28 °C on either trypticase soya agar (TSA; Oxoid), in trypticase soya broth (TSB; Oxoid), in potato dextrose broth (PDB; Biocorp) or in M9 minimal medium (MP Biomedicals) supplemented with glucose (Sigma-Aldrich) to a final concentration of 0.4%. Liquid cultures were agitated during incubation (120 rpm). To solidify the media, 15 g mL^-1^ bacteriological agar (Oxoid) was added. STA medium (per 1 L: 30 g trypticase soya broth (Oxoid) and 7 g bacteriological agar (Oxoid)) was used as the soft top agar. When required, the growth media were supplemented with various concentrations of mitomycin C (Abcam), chloramphenicol (A&A Biotechnology), ampicillin (A&A Biotechnology), ciprofloxacin (Sigma-Aldrich), norfloxacin (Abcam) or hydrogen peroxide (Laboratorium Galenowe Olsztyn, Poland) as described below.

### Induction, purification, and estimation of tailocin titer after induction

P2D1 tailocins were induced, purified, concentrated, and tittered as described in ^10^. Briefly, *Dickeya dadantii* strain 3937 was grown overnight (ca. 16 h) in TSB at 28°C with shaking (120 rpm). The cultures were then rejuvenated by diluting them 1:40 in 10 or 100 mL of fresh TSB medium. The diluted culture grew for 2.5 hours under the same conditions. Such prepared *D. dadantii* cultures were then supplemented with mitomycin C (Abcam, Poland) to a final concentration of 1 μg mL^-1^ to induce the production of P2D1 tailocins. Following mitomycin C treatment, the cultures were incubated for another 24 h at 28 °C with shaking (120 rpm). 2.2 mL of each post-induction bacterial culture was cleared from bacterial cells through centrifugation (10 min., 8000 RCF, 22 °C). The resulting supernatant was filtered (0.2 µm, PES membrane, Googlab), and 2 mL of the filtrate was transferred to tubes containing polyethylene glycol (PEG-8000, Promega) to achieve the final PEG concentration of 10%. The samples were incubated for 16-20 hours at 4 °C with gentle horizontal shaking to precipitate tailocins. The precipitant was harvested by centrifugation (1 h, 16000 RCF, 4 °C). After removing the supernatant, the samples were dried under laminar flow until the remaining liquid evaporated. The resulting tailocins were resuspended in 200 μL of phosphate-buffered saline (PBS, pH 7.2), yielding a tenfold concentration relative to the initial volume. The purified tailocins were stored at 4 °C until further analysis.

The concentration of tailocin particles was determined using a semi-quantitative spot test assay, as described earlier ^19^. Briefly, Petri dishes containing TSA were overlaid with soft top agar (STA). Before pouring, the soft top agar was cooled to 48 °C and inoculated (1:60) with an overnight culture of tailocin-sensitive strain *M. paradisiaca* NCPPB 2511 ^10^. The samples tested for tailocin titer were serially twofold diluted in PBS (pH 7.2). Then, in duplicates, 2 μL of each tested suspension was spotted on the plates inoculated with *M. paradisiaca*. After overnight incubation at 28 °C, the highest dilution of tailocins causing the formation of clear zones (plaques) on the bacterial lawn was determined. Since the tested method for determining tailocin concentration is based on assessing their activity, the results were expressed as relative activity in arbitrary units (AU), with 1 AU being defined as equal to the highest dilution that still caused a visible plaque on the lawn of sensitive bacterial strain (*Musicola paradisiaca* strain NCPPB 2511).

### Effect of mitomycin C concentration on P2D1 yield

To determine whether the concentration of mitomycin C affects the induction and the final yield of the P2D1 tailocins in *D. dadantii* strain 3937, we tested eleven concentrations of mitomycin C. *D. dadantii* strain 3937 was grown for 16 hours in 10 mL of TSB at 28 °C with shaking (120 rpm). The cultures were then diluted 1:40 in fresh TSB, incubated under the same conditions for 2.5 hours, and then spiked with 0.1, 0.2, 0.3, 0.4, 0.5, 1.0, 1.5, 2.0, 3.0, 4.0, or 10 µg mL^-1^ of mitomycin C (Table S1). Following the addition of the inducer, the resulting bacterial cultures were further incubated at 28 °C for 6 h with shaking (120 rpm). In each case, tailocin titer was estimated as described above. The experiment was independently conducted four times with two replicates each, and the results were averaged.

### Effect of mitomycin C on the viability of *D. dadantii* 3937 cells

*D. dadantii* strain 3937 was cultured in TSB medium for 16 hours at 28 °C with shaking (120 rpm). The overnight cultures were rejuvenated (1:40) in 250 mL of fresh TSB medium, and the incubation was continued under the same conditions for 2.5 hours. Subsequently, mitomycin C was added to a final concentration of 1 µg mL^-1^, and the cultures were continued under the same conditions. Samples were collected at various time points during the incubation: 2.5 hours before mitomycin C addition, at the time of inducer administration (T=0), and at 0.5, 1, 2, 4, 6, 8, and 24 hours post-treatment. The control sample consisted of bacterial cultures not subjected to mitomycin C treatment. To obtain cell count, all harvested samples were serially diluted tenfold in PBS (Sigma-Aldrich, pH 7.2), and 10 µl aliquots of the dilutions were plated on TSB agar plates by spotting as described earlier ^20^. The plates were dried under a laminar flow until the liquid was fully absorbed and incubated at 28 °C for 20 hours. Following incubation, the number of colonies was counted, and the average colony-forming units (CFU) per milliliter of the original culture were calculated. The experiment used two biological replicates, each containing three technical ones.

### Yield of P2D1 tailocins at different time points after induction

To examine the changes in P2D1 concentration following the addition of the inducing agent, strain *D. dadantii* 3937 was grown for 16 hours in 10 mL of TSB at 28 °C with shaking (120 rpm). The cultures were then diluted 1:40 in fresh TSB and incubated under the same conditions for an additional 2.5 hours. To induce the production of tailocins, bacterial cultures were spiked with 1 µg mL^-1^ of mitomycin C, and the resulting culture was incubated under the same conditions for another 24 h. Culture samples were collected at T=0 (addition of mitomycin C to *D. dadantii* strain 3937 culture) and after 0.5, 1, 2, 4, 6, 8, and 24 h after induction. In each case, tailocin titer was estimated as described above using the semi-quantitative spot test with *M. paradisiaca* NCPPB 2511 as a reporter strain. The experiment was independently repeated three times, and the results were averaged.

### Expression of genes involved in P2D1 synthesis with RT-qPCR

To assess the expression of structural genes encoding P2D1, the expression of genes coding for fiber, tube, and sheath, as well as the reference genes *lpxC* and *rplU,* was analyzed using RT-qPCR.

#### Isolation of RNA

Total RNA from bacterial cultures was isolated using the RNeasy Protect Bacteria Mini Kit (QIAGEN) according to the manufacturer’s protocol involving proteinase K treatment. The processed samples included the cultures of *D. dadantii* 3937 grown as described for determining the activity of tailocin inducers, harvested immediately before the addition of mitomycin C (T=0) and at 1, 2, and 4 hours after treatment. Samples untreated with mitomycin C were processed the same way as a control. For time points 0, 1, and 2 h post-induction, 5 ml of the culture was mixed with RNAprotect Bacteria Reagent (QIAGEN) at a 1:2 ratio, and the cells were pelleted (10 min., 5000 RCF, 22 °C) for isolation of RNA. For mitomycin-treated samples collected after 4 hours, due to the loss in viable cell count, a larger volume (50 ml) of the culture was harvested, pelleted (3 minutes, 8000 RCF, 22 °C), resuspended in 500 μl of PBS buffer, and immediately mixed with 1 ml of the RNAprotect Bacteria Reagent. The increased culture volume processed for timepoint T=4 h did not apply to the control (untreated) samples. The purified RNA was treated with the TURBO DNA-free Kit (Thermo Fisher Scientific) to remove potential genomic DNA contamination. The integrity of RNA was verified by agarose gel electrophoresis, and its concentration was determined using NanoDrop 2000 (Thermo Fisher Scientific). The samples were stored at −80 °C for further use.

#### cDNA synthesis and real-time qPCR

RNA was reverse transcribed into cDNA using the Transcriptor First Strand cDNA Synthesis Kit (Roche) with random hexamer primers and with the optional denaturation step. For each reaction, 500 ng of RNA was used as a template. Real-time qPCR was performed on the CFX96 instrument (Bio-Rad) using Power SYBR Green PCR Master Mix (Thermo Fisher Scientific), as described before ^21^. The template cDNA was diluted 1:4. All primers used in the experiment were designed using Primer3Plus ^22^ (Table S2). Primer specificity was verified by agarose gel electrophoresis and melt curve analysis. Primer efficiency was established based on serial dilutions of post-PCR products as a template. Gene expression analysis was performed using CFX Maestro 2.3 software (Bio-Rad). The expression of target genes was normalized to *lpxC* and *rplU* genes, which were previously established as valid reference genes in *D. dadantii* ^23^. Gene expression was scaled to the reference biological group, which was the non-treated sample collected at 0 hours (T=0), corresponding to the time of mitomycin C addition to the experimental group.

### Effect of antibiotics on P2D1 yield

To test whether chemicals other than mitomycin C can induce the production of P2D1 tailocins in *D. dadantii* strain 3937, four antibiotics with different modes of action were selected and used in a model experiment: chloramphenicol – inhibiting protein synthesis, ampicillin – that inhibits bacterial cell wall synthesis, ciprofloxacin, and norfloxacin – both inhibiting DNA replication and repair. *D. dadantii* strain 3937 was grown for 16 hours in 10 mL of TSB at 28 °C with shaking (120 rpm). The cultures were then diluted 1:40 in fresh TSB and incubated under the same conditions for an additional 2.5 hours. To induce the production of tailocins, bacterial cultures were spiked with either chloramphenicol (final concentration: 4 µg mL^-1^), ampicillin (final concentration: 0.004 µg mL^-1^), ciprofloxacin (final concentration: 0.016 µg mL^-1^) and norfloxacin (final concentration: 0.016 µg mL^-1^)(Table S1). As a positive control and known P2D1 inducer, mitomycin C (final concentration: 1 µg mL^-1^) was used. After adding putative inducers, the resulting bacterial cultures were further incubated at 28 °C for 6 h with shaking (120 rpm). In each case, tailocin titer was estimated as described above using the semi-quantitative spot test with *M. paradisiaca* NCPPB 2511as a reporter strain. The obtained results were expressed as relative activity in arbitrary units (AU) as described above. The experiment was independently conducted four times, containing two technical replicates each, and the results were averaged.

### Effect of hydrogen peroxide on P2D1 yield

To assess the effect of hydrogen peroxide on tailocin induction in *D. dadantii* 3937 and to compare it with the bacterial cell response induced by mitomycin C, an experiment was conducted following similar conditions as those described above to test the effect of varying concentrations of mitomycin C. The only difference in treatment was that four concentrations of hydrogen peroxide (0.5, 1, 5, and 10 mM) were tested instead of spiking cultures with mitomycin C. A single concentration of mitomycin C (1 µg mL^-1^) and an untreated bacterial culture were positive and negative controls, respectively. The yield of P2D1 particles obtained following each treatment was expressed in relative units (AU) as described above. The experiment was conducted three times, each time with two technical replicates.

### Effect of the type of growth medium on P2D1 yield

To test whether the type and composition of the growth medium affect the production of P2D1 tailocins, three different media were tested: M9 minimal medium with 0.4% glucose, PDB (rich medium with potato extract and glucose), and TSB (rich medium with peptone and glucose). *D. dadantii* strain 3937 was grown for 16 hours in 10 mL of TSB at 28 °C with shaking (120 rpm). The cultures were diluted 1:40 in fresh TSB, PDB, or M9+0.4% glucose medium and incubated under the same conditions for 2.5 hours. To induce the production of tailocins, bacterial cultures were spiked with 1.0 µg mL^-1^ of mitomycin C. Following the addition of the inducer, the resulting bacterial cultures were further incubated at 28 °C for 6 h with shaking (120 rpm). In each case, tailocin titer was estimated as described above using the semi-quantitative spot test with *M. paradisiaca* NCPPB 2511 as a reporter strain. The experiment was independently repeated three times, with two replicates each, and the results were averaged.

## Results

### P2D1 output depends on the concentration of mitomycin C

*D. dadantii* 3937 was reported to produce P2D1 dickeyocins at a baseline level of ca. 6.5 log_2_AU when grown in TSB in the absence of an inducer. However, the production was upregulated following treatment with 0.1 µg mL^-1^ of mitomycin C ^10^. Here, we established the inductive effect of mitomycin C across various concentrations (0.1 – 10 µg mL^-1^). Although mitomycin C significantly upregulated P2D1 production at all concentrations tested, the highest yield was observed for concentrations between 0.5 and 1.5 µg mL^-1^. At these concentrations, activity levels approached 12.5 log_2_AU, indicating approximately a 64-fold increase relative to the untreated control sample (Fig. 1 A).

**Figure 1.**
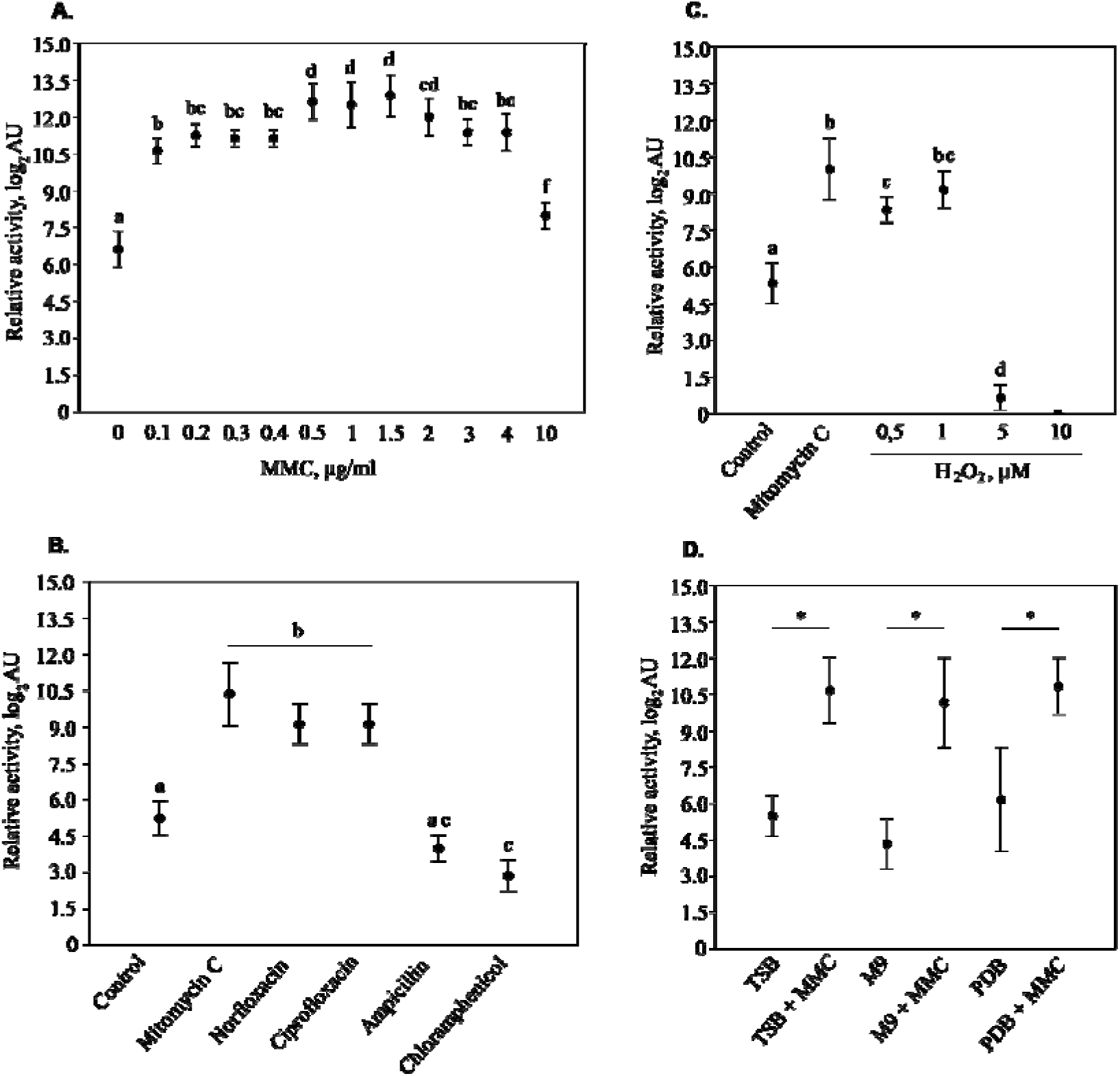
Production of tailocins by *D. dadantii* 3937 depending on the applied concentration of mitomycin C, the growth medium, and the type of inducer. Tailocin titer was expressed as relative tailocin activity in arbitrary units (AU), with 1 AU defined as the reciprocal of the highest dilution that caused a visible plaque on a lawn of susceptible strain *M. paradisiaca* NCPPB 2511. In all panels, data points show mean values, and error bars represent standard deviations. Statistically significant differences between groups for analyses in panels A and C were determined using Tukey’s pairwise comparison ^41^, followed by Copenhaver Holland post hoc analysis ^42^, while for analyses in panels B and D, these were determined by Kruskal-Wallis test ^43^, followed by Dunn’s post hoc test ^44^. In all panels, groups labeled with the same letter are not significantly different (α=0.05). Panel (**A**) depicts the titer of P2D1 tailocins depending on the applied concentration of mitomycin C (MMC). Panel (**B**) shows tailocin titer when testing the inducive potential of selected antibiotics: norfloxacin 0.016 µg mL^-1^; ciprofloxacin 0.016 µg mL^-1^; ampicillin 0.004 µg mL^-1^; chloramphenicol 4 µg mL^-1^. Treatment with mitomycin C (1 µg mL^-1^) was used as a reference. Control – culture with no potential inducer added. Panel (**C**) shows tailocin yield following induction with different concentrations of hydrogen peroxide: 0.5, 1, 5, and 10 mM, with mitomycin treatment as reference (1 µg mL^-1^). **Control** – culture with no potential inducer added. Panel (**D**) shows tailocin yield in different growth media when induced with mitomycin C (MMC, 1 µg mL^-1^): TSB – Trypticase Soya Broth; M9 – M9 minimal medium with 0.4% glucose; PDB – Potato Dextrose Broth

### Mitomycin C causes a decline in viable *D. dadantii* 3937 cell count that is caused by the accumulation of P2D1 tailocins

The impact of mitomycin C (1 µg mL^-1^) on the growth and survival of *D. dadantii* strain 3937 in TSB medium was evaluated. In the control culture, which was not exposed to mitomycin C, a robust 34-fold increase in viable bacterial cells was observed over the 24-hour incubation period, with the count rising from 7.7 to 9.2 log_10_CFU mL^-1^ (Fig. 2 A). However, in the presence of mitomycin C, bacterial survival dropped sharply. Four hours after the addition of the inducer, the number of viable cells decreased approximately 800-fold, from 7.6 to 4.7 log_10_CFU mL^-1^. This decline became even more pronounced at 6 hours, where the cell count fell by 3.6 million-fold to just 1 log_10_ CFU mL^-1^. By the end of the 24-hour incubation with mitomycin C, the bacterial population was nearly eradicated, with a final count of 0.4 log_₁₀_ CFU mL^-1^.

**Figure 2.**
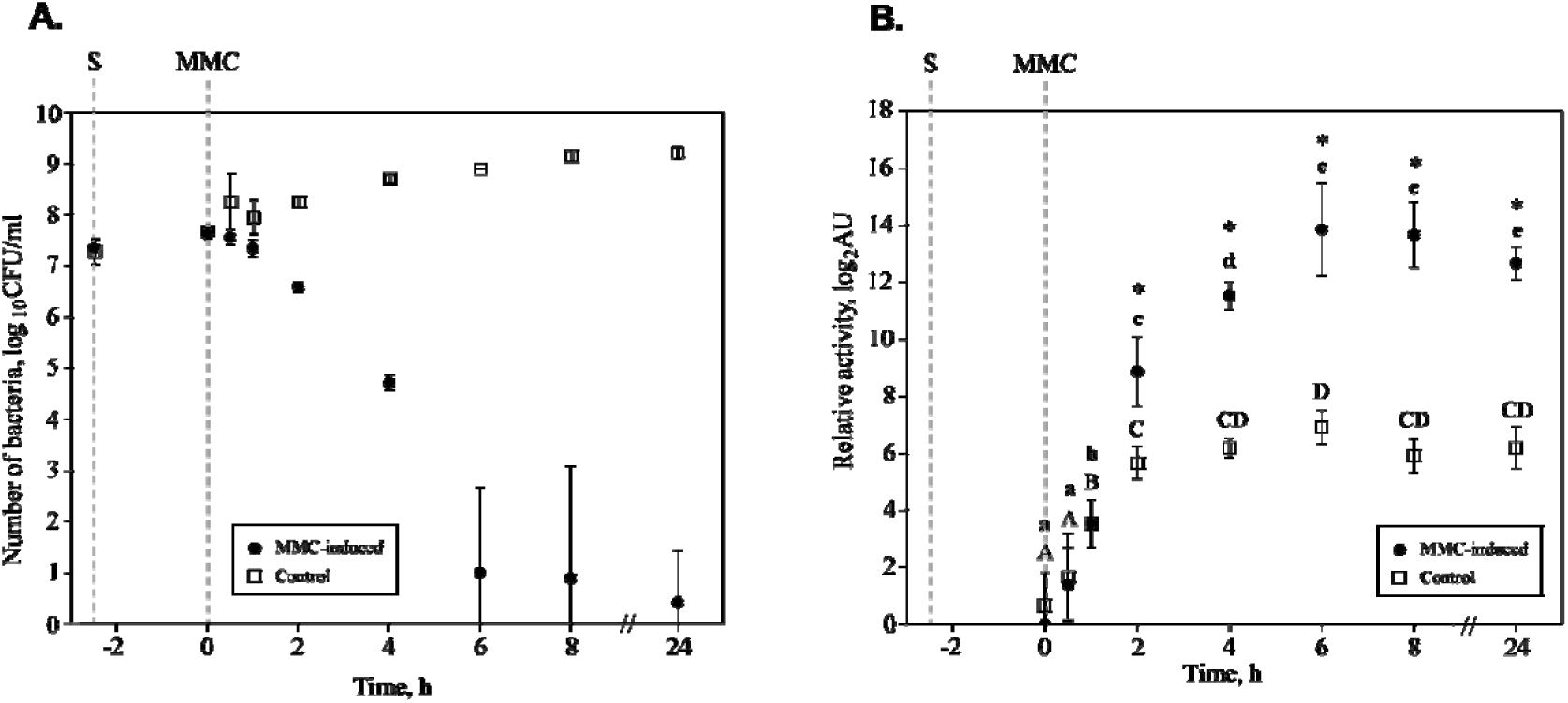
Tailocin yield and viable bacterial cell count as a function of time following the addition of inducer. (**A**) Survival of *D. dadantii* strain 3937 over time, measured in colony-forming units (CFU). Data points represent the viable bacterial cell count in cultures treated with mitomycin C at a concentration of 1 µg mL^-1^. The counts are shown at various intervals, illustrating the decline in bacterial viability as the cells produce and release tailocins.(**B**) The titer of P2D1 tailocins isolated from the culture of *D. dadantii* strain 3937 at different time points (0-24h) post induction with mitomycin C (1 µg mL^-1^). The x-axis indicates the incubation time of bacterial cells with the inducer, after which tailocins were harvested for activity against *M. paradisiaca* NCPPB 2511. Cultures unexposed to mitomycin C were used as controls. Points represent the mean activity (mean AU) of measurements from 3 independent experiments. Error bars indicate standard deviation ranges. S represents the start of the culture, and MMC indicates the time of spiking the culture with mitomycin C. In panel (B), significant differences between MMC-treated samples collected at different time points were determined using the Mann-Whitney U test ^45^. Groups labeled with the same letter are not significantly different (α=0.05). Small letters are used for MMC-treated samples and capital letters for control. **Asterisks (*) indicate statistically significant differences (p < 0.05; Mann-Whitney U test** ^45^ **between experimental and control samples collected at the same time points.**

### P2D1 yield reaches maximum level six hours after induction with mitomycin C

Statistically significant differences between the mitomycin C-treated samples and the untreated control were detectable 2 hours post-induction, with the bactericidal activity of the induced tailocins increasing 9-fold (to 8.86 log_2_AU) relative to the control (5.67 log_2_AU) (Fig. 2 B). The highest tailocin titer was observed 6 hours after mitomycin C addition, with an average value of 13.9 log_2_AU, representing an approximately 123-fold increase compared to the untreated control. No statistically significant differences in P2D1 yield were observed between mitomycin C-treated samples collected at 6 to 24 hours post-induction, indicating that after reaching its maximum, the titer of P2D1 particles remained stable (Fig. 2 B).

### Expression of genes encoding P2D1 structural genes peaks two hours after induction

The expression of three genes encoding structural proteins of dickeyocin P2D1 (responsible for forming the sheath, tube, and tail fiber) was analyzed using RT-qPCR to monitor the tailocin gene expression. Gene expression was compared between *D. dadantii* strain 3937 cells treated with mitomycin C (1 µg mL^-1^) and untreated control cells. RNA samples were collected at four time points following mitomycin C addition (0, 1, 2, and 4 h after induction). Gene expression levels were independently normalized to the control samples collected at 0 h for each gene. In the control group (no inducer added), no significant changes in gene expression were observed over time, except for a slight increase in the expression of the sheath protein gene at 4 hours (Fig. 3). In contrast, cells exposed to mitomycin C showed a marked increase in the expression of all target genes. After 1 hour, the expression of the genes encoding the sheath, tube, and fiber proteins increased by approximately 15-, 9-, and 4-fold, respectively. The peak expression was recorded 2 hours post-induction, with the tube protein gene showing a 655-fold increase and the sheath and fiber genes increasing by 295- and 191-fold, respectively, compared to the control (Fig. 3, Table S3). By 4 hours, a declining trend was observed for the expression of all three tailocin genes, with the difference being statistically significant for the fiber and sheath genes (Fig. 3).

**Figure 3.**
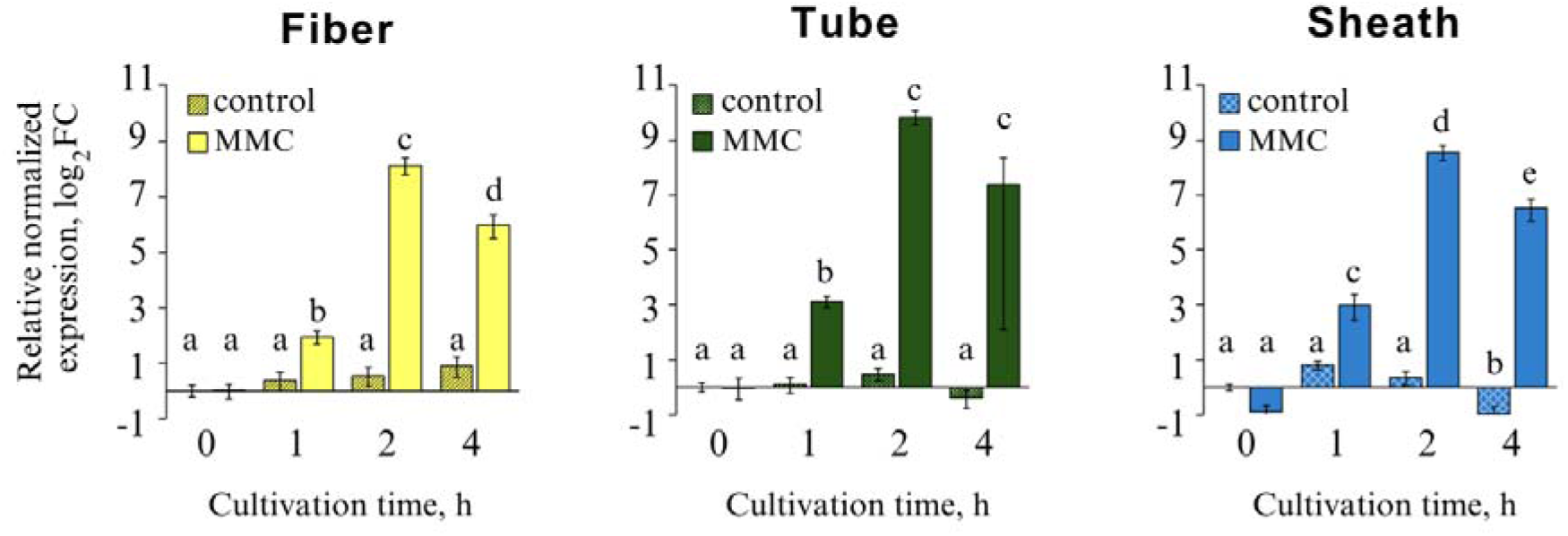
Expression of P2D1-encoding structural genes in *D. dadantii* 3937 at different time points following induction with mitomycin C. The graphs depict the relative expression levels of three target genes: fiber (DDA3937_RS12070), tube (DDA3937_RS12115), and sheath (DDA3937_RS12110), in samples collected after specified incubation times following the addition of mitomycin C (1 µg mL^-1^). Control samples consist of bacteria that were not treated with the antibiotic. The fold change (log2) in gene expression for the target genes was calculated relative to the control samples collected at time 0. Statistically significant differences among the experimental groups were determined using one-way ANOVA ^46^, followed by Tukey’s Honest Significant Difference post hoc analysis ^47^. Groups marked with the same letter do not differ significantly (α=0.05).

### P2D1 tailocins are induced by antibiotics, affecting the replication and repair of bacterial genomic DNA

We examined the impact of selected antibiotics (viz. ampicillin, ciprofloxacin, norfloxacin, and chloramphenicol) on the output of dickeyocin P2D1. Notably, norfloxacin and ciprofloxacin significantly increased tailocin production, with tailocin activity levels rising 14-fold compared to the control (untreated culture). These results were statistically comparable to the induction of P2D1 with mitomycin C. In contrast, chloramphenicol caused a five-fold decrease in P2D1 production, whereas ampicillin showed no significant differences in P2D1 output relative to the control (untreated cultures) (Fig. 1 B).

### P2D1 tailocins are inducted by hydrogen peroxide

Supplementation of *D. dadantii* 3937 culture with hydrogen peroxide significantly affected dickeyocin P2D1 production, as evidenced by changes in bactericidal activity against *M. paradisiaca* NCPPB 2511 (Fig. 1 C). The highest mean induction was observed by adding 1 mM H_2_O_2_ to the culture, increasing bactericidal activity to an average of 9.2 log_2_AU, making it 14.3 times higher than the control (5.3 log_2_AU). The effect of 1 mM H_2_O_2_ was not significantly different from that of the reference treatment with 1 µg mL^-1^ of mitomycin C. H_2_O_2_ at 0.5 mM increased activity 8-fold (to 8.3 log_2_AU). In contrast, activity following treatment with the highest tested concentrations, 5 and 10 mM H_2_O_₂,_ was negligible (Fig. 1 C).

### The type of the growth medium affects neither the basal level nor the induction of P2D1 tailocins

We found that the type of microbiological medium used during the cultivation of the *D. dadantii* 3937 strain did not affect the level of production of dickeyocin P2D1 (mean 5.3 log_2_AU). This was true both for the baseline level of tailocin production as well as following induction with mitomycin C (mean 10.6 log_2_AU) (Fig. 1 D), as no statistically significant differences (α=0.05) were observed within the untreated and treated groups, irrespective of the growth medium.

## Discussion

Although multiple studies have investigated the genetic factors associated with the production of phage tail-like particles (tailocins), wide-ranging analyses examining altogether the interplay between tailocin inducers, induction timing, structural gene expression, and the viability of producing bacterial cells remain scarce. This study is the first to document the sequential dynamics of the production of P2D1 tailocin in plant pathogenic *Dickeya* spp. ^24^ and specifically in a model strain *D. dadantii* 3937 ^16^ (see Figure 4 for a summary chart). Likewise, we analyzed the ecological perspective of P2D1 production and how this process can be triggered under environmental conditions and *via* ecological factors.

**Figure 4.**
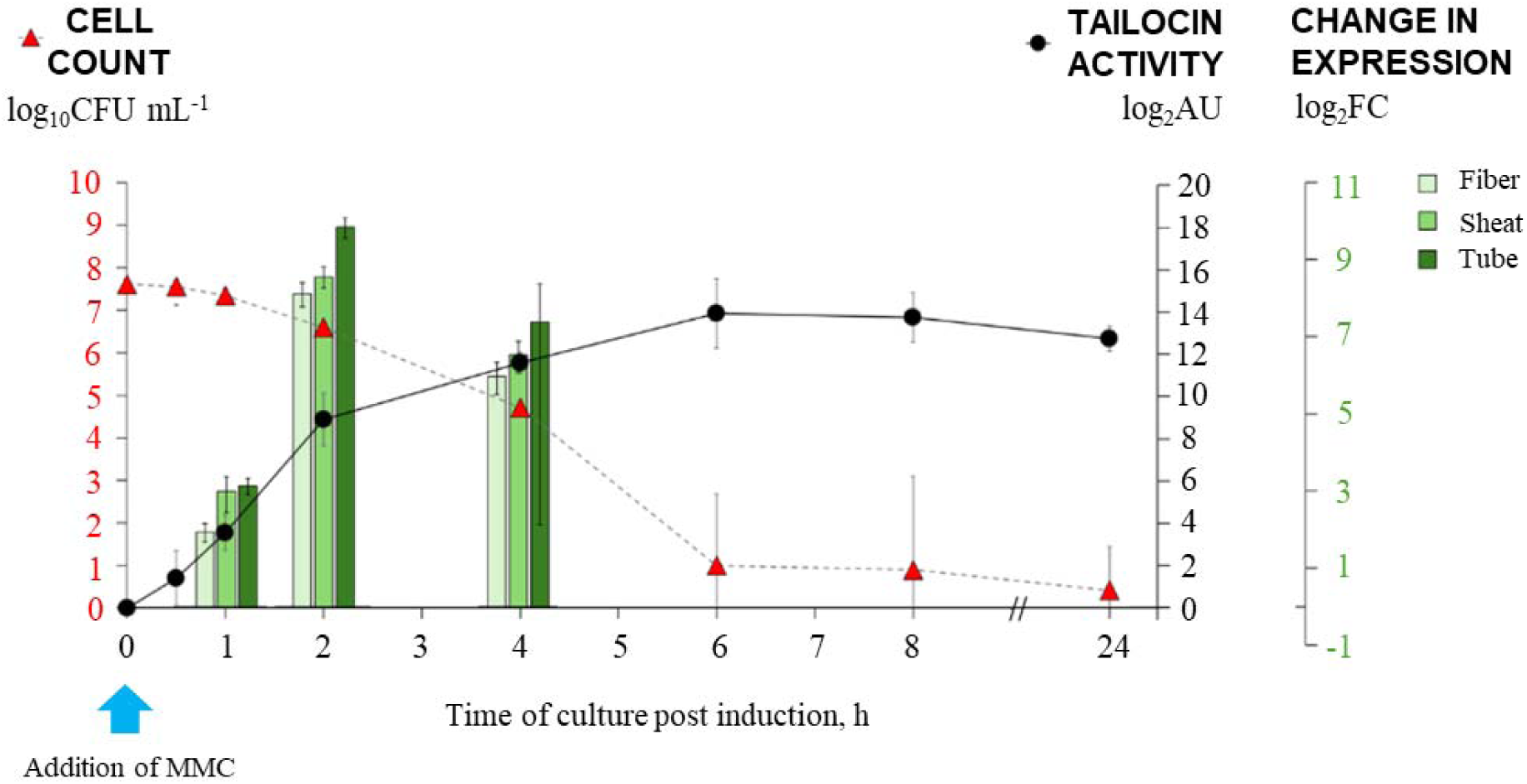
Temporal response of *D. dadantii* 3937 to mitomycin C. The figure illustrates the dynamics of strain 3937 in response to stress induced by mitomycin C. The graph presents the number of viable bacterial cells producing P2D1 tailocins, the relative activity of the produced tailocins against *M. paradisiaca* NCPPB 2511, and the relative expression levels of genes encoding structural proteins of the tailocins. The cultures were treated with mitomycin C (MMC) at a concentration of 1 µg mL^-1^, with the inducer added to the bacterial culture at time 0.

Our study demonstrated that the production of P2D1 tailocins is significantly influenced by mitomycin C. This observation confirmed its important role as a potent phage tail-like particle inducer in *D. dadantii* 3937, which we have demonstrated earlier ^10^. It also extends these findings by characterizing the dose-dependent effects of mitomycin C. The marked upregulation of P2D1 production was observed across a range of inducer concentrations. Our observations align with findings on other bacteriocins, where moderate stress levels maximize bacteriocin synthesis ^25^. However, the highest (above 1.5 µg mL^-1^) concentrations of mitomycin C used in our study resulted in significantly decreased P2D1 yields. This is consistent with studies concerning the impact of mitomycin on cells, where excessive DNA damage due to mitomycin C treatment overwhelms cellular repair machinery, resulting in impaired cellular function or even death ^26^, therefore reducing tailocin yield.

The induction of P2D1 tailocins with mitomycin C caused a remarkable decline in viable *D. dadantii* cells (ca. 3.5-million-fold reduction observed just after 6 h post-induction). Such an observation aligns well with the known canonical mechanism of tailocins production, where the synthesis and release of these particles disrupt the cell membranes and/or negatively affect other cellular processes, resulting in the rapid death of the producers ^4,27^. Furthermore, stabilizing tailocin titers beyond six hours after induction suggests that their production may cease once a certain threshold is reached. This could be attributed to the existence of a limit on the number of cells within the population capable of producing tailocins, driven by their heterogeneous response to stress factors ^28^. This heterogeneous response means that not all cells activate tailocin production, resulting in a subset of cells that do not undergo lysis and thus survive. As to our knowledge, at the moment, there are no similar studies evaluating the production of phage tail-like particles in different bacterial species; we can only speculate that our observations may mirror the situation observed in colicin production, where a self-limiting loop of regulation ensures efficient resource allocation, during stress ^29^.

Real-time analysis of the temporal expression of P2D1 structural genes revealed that peak transcription occurred two hours after induction with a significant upregulation of genes encoding P2D1 tube, fiber, and sheath. This fast transcriptional response is rather characteristic of bacteriocin systems regulated by the SOS response ^30^, where it triggers the expression of genes inducible by the DNA damaging agents ^31,32^. Similar temporal patterns have been reported in other bacterial species, where gene expression peaks early in the stress response, followed by a decline as the cells transit into a survival state ^6,33^. The declining trend observed after 4 hours may suggest that transcriptional activation of tailocin genes is tightly regulated. Such a synchronized response ensures that the synthesis of tailocins, including P2D1, does not compromise cellular integrity during prolonged stress ^34^. In other microorganisms, the regulation of tailocin synthesis involves complex mechanisms that balance positive and negative regulatory factors. For instance, in *Pseudomonas aeruginosa*, tailocin production is controlled by the positive regulator PrtN and the negative regulator PrtR, with additional oversight provided by the LexA protein, which acts as a gatekeeper for the SOS response ^35^. In *Stenotrophomonas maltophilia*, MpsA and MpsH function as positive regulators, while MpsR acts as a repressor under normal conditions, preventing unnecessary tailocin production ^36^. The proteins and mechanisms involved in regulating the production of P2D1 tailocin in *Dickeya dadantii* 3937 and the buffering system that allows a portion of the population to survive despite the high cost of tailocin production under inducing conditions remain to be elucidated.

The induction of P2D1 by chemicals other than mitomycin C (i.e., antibiotics and hydrogen peroxide) provides valuable insights into the versatility of tailocin regulation, including processes regulating P2D1 synthesis in *D. dadantii* strain 3937. Not surprisingly, antibiotics such as ciprofloxacin and norfloxacin, which inhibit bacterial replication, may mimic the effect of mitomycin C by inducing the SOS response, as seen in other studies ^37,38^. Similarly, oxidative stress caused by hydrogen peroxide significantly upregulated tailocin production in strain 3937 at a comparable level to mitomycin C. The latter observation supports the hypothesis that tailocin synthesis is under the control of the generalized stress response ^4,9,27^. However, the neglectable production of P2D1 tailocins at higher hydrogen peroxide concentrations highlights the importance of stress insensitivity in modulating tailocin output ^39^. In contrast, as expected, chloramphenicol inhibited P2D1 production, likely by impairing protein synthesis and interference with cell wall synthesis, respectively ^40^.

The lack of significant differences in P2D1 yield across several growth media highlights the robustness of tailocin synthesis in *D. dadantii* strain 3937. These findings may contrast with studies on other phage tail-like particles, where medium composition heavily influenced tailocin yield due to the differences in nutrient availability or pH ^6,19^. The medium independence of P2D1 production suggests that tailocin regulation is predominantly governed by cellular stress pathways rather than other factors.

To conclude, our study gives new insights into the events during the assembly and release of P2D1 in *D. dadantii* strain 3937 and broader into the regulation of tailocin synthesis in *Dickeya* spp. and other closely related plant pathogenic bacteria. To our knowledge, this is the first investigation to provide detailed insights into the sequential events of tailocin assembly and release under various inducing conditions. The lack of comparable studies highlights the importance of further exploration into tailocin regulation and its implications for bacterial stress responses and interspecies competition.

## Data availability

Data generated or analyzed during this study are included in this article (including its Supplementary Information files). These data can also be obtained from the corresponding author and shared freely upon reasonable request. Correspondence and requests for materials should be addressed to Robert Czajkowski.

## Institutional Review Board statement

Not applicable

## Insitutuonal concent statement

Not applicable

## Author contributions

MS: Conceptualization, Investigation, Methodology, Visualization, Writing – original draft, Writing – review & editing, DMK: Conceptualization, Investigation, Supervision, Visualization, Writing – original draft, Writing – review & editing, MB: Investigation, Methodology, Writing – original draft, Writing – review & editing, RC: Conceptualization, Data curation, Funding acquisition, Resources, Supervision, Writing – original draft, Writing – review & editing

## Acknowledgments

This research was financially supported by the National Science Center, Poland (Narodowe Centrum Nauki, Polska) via a research grant SONATA BIS 10 (2020/38/E/NZ9/00007) to Robert Czajkowski.

## Supplementary Figures

**Figure S1.**
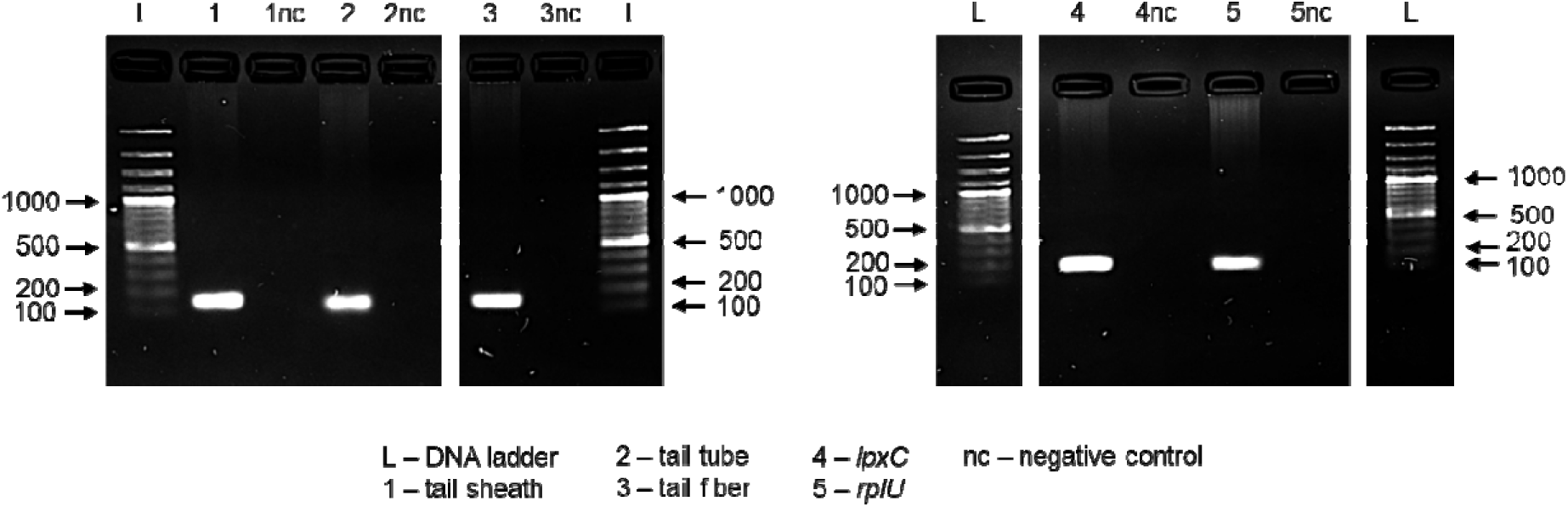
Specificity of primers applied in real-time qPCR. Agarose gel electrophoresis (2%) depicting single PCR products of the expected sizes for each of the analyzed target genes. L – GeneRuler™ 100bp DNA Ladder Plus (Thermo Fisher Scientific). nc – no template negative control; DNA staining was done using Novel Juice (GeneDireX). The image was captured with a ChemiDoc XSR (Bio-Rad) and processed using Image Lab software.

**Figure S2.**
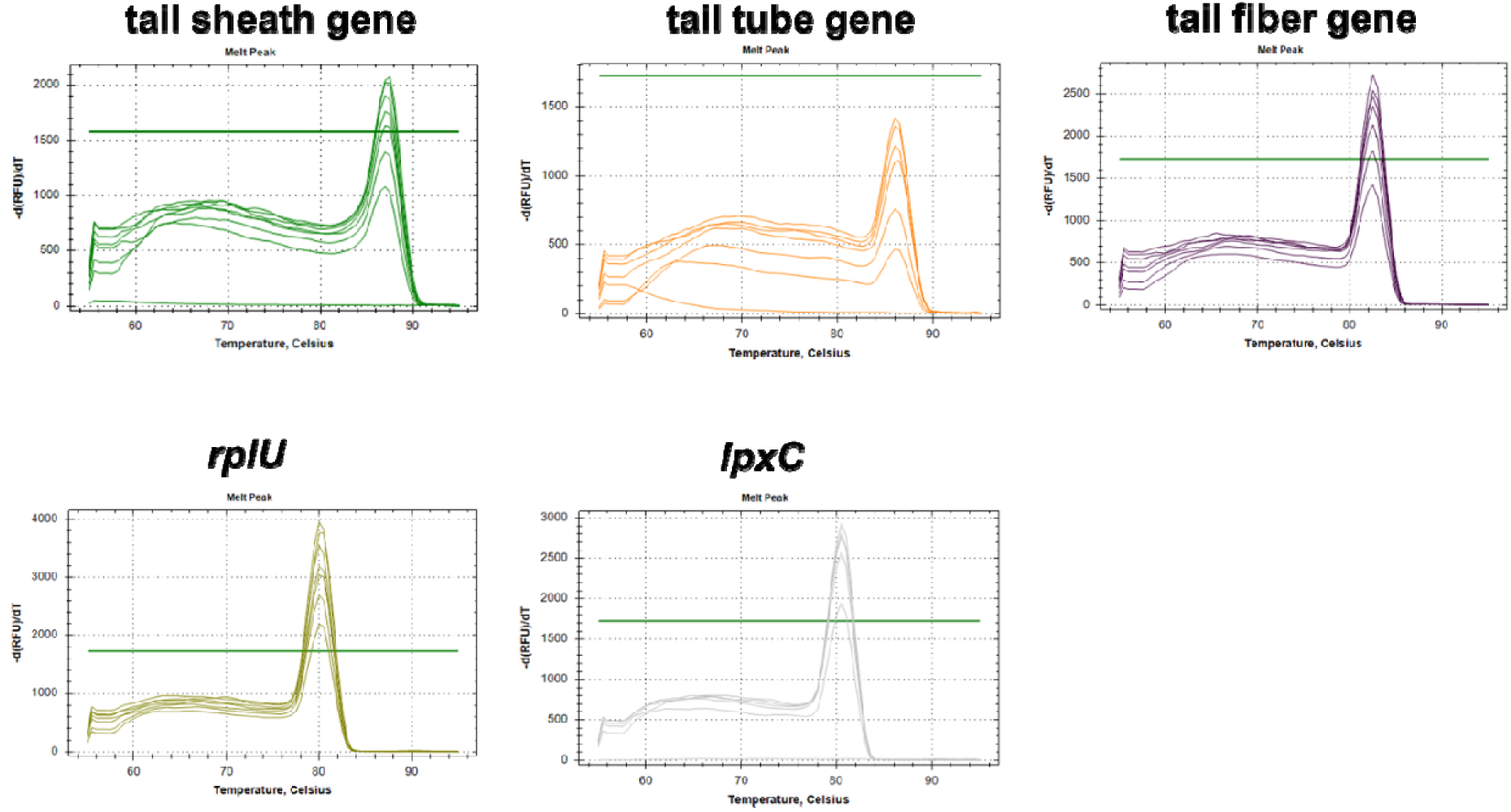
Melt curve analysis. Melt curve analysis (55-95 °C with 0.5 °C increment every 5 seconds) for the amplicons of each target gene, demonstrating that each pair of PCR primers produces a single peak. The gene loci corresponding to the designated targets are provided in Table S2.

**Figure S3.**
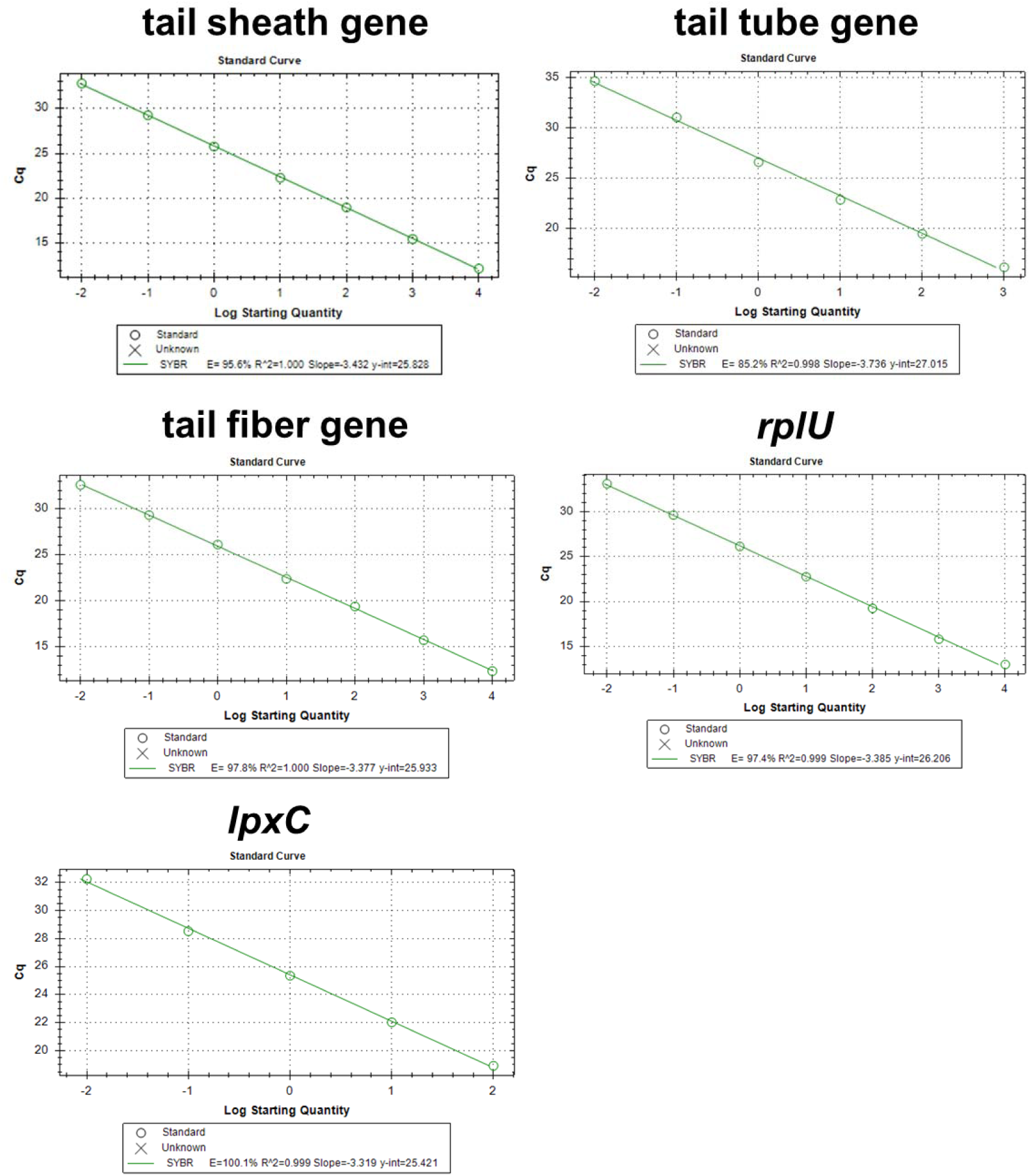
Standard curves for the estimation of PCR efficiency. Standard curves were used to determine PCR efficiency for the reference genes (*rplU* and *lpxC*) and the target genes (tail sheath gene, tail tube gene, and tail fiber gene). The gene loci corresponding to the designated targets are provided in Table S2.

## Supplementary Tables

**Table S1.** Conditions for the induction of tailocin production.

| Experiment | Inducer | Concentration | Unit | Incubation time, h |
| --- | --- | --- | --- | --- |
| Effect of mitomycin C concentration on tailocin yield | Mitomycin C | 0.1; 0.2; 0.3; 0.4; 0.5; 1; 1.5; 2; 3; 4; 10 | $\mu\text{g mL}^{-1}$ | 24 |
| Tailocin yield at different time points after induction | Mitomycin C | 1 | $\mu\text{g mL}^{-1}$ | 0; 0.5; 1; 2; 4; 6; 8; 24 |
| Potential of selected inducers to induce tailocins | Mitomycin C | 1 | $\mu\text{g mL}^{-1}$ | 6 |
| | Chloramphenicol | 4 | $\mu\text{g mL}^{-1}$ | 6 |
| | Ampicillin | 0.004 | $\mu\text{g mL}^{-1}$ | 6 |
| | Ciprofloxacin | 0.016 | $\mu\text{g mL}^{-1}$ | 6 |
| | Norfloxacin | 0.016 | $\mu\text{g mL}^{-1}$ | 6 |
|  | Hydrogen peroxide | 0.5; 1; 5; 10 | mM | 6 |

**Table S2.**
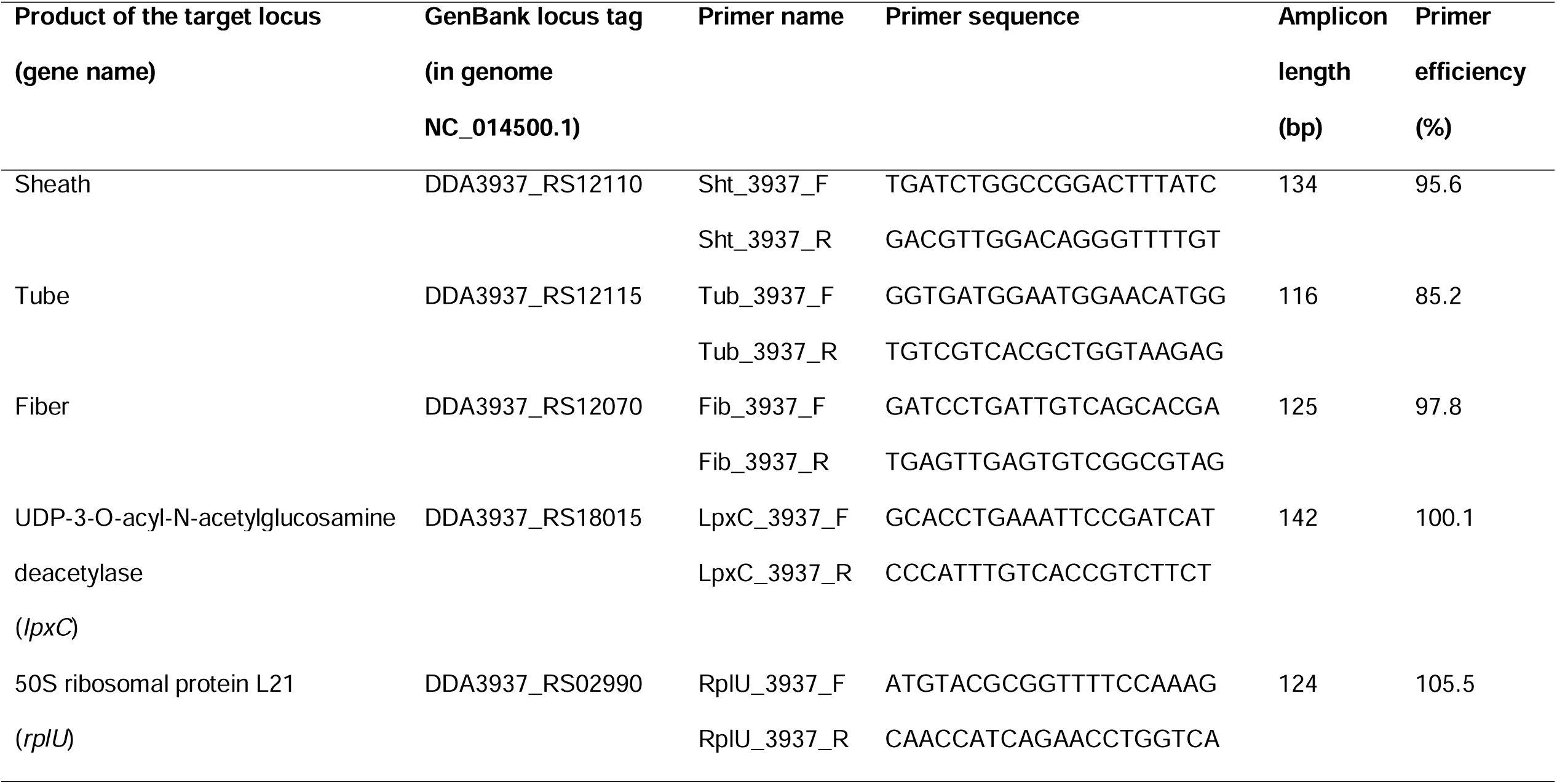
Primers designed and used in this study.

**Table S3.** Expression of structural genes encoding P2D1 at different time points following induction.

| Gene<br>(locus<sup>A</sup>) | time<br>(h) | Control |  |  | Mitomycin C treatment |  |  |
| --- | --- | --- | --- | --- | --- | --- | --- |
|  |  | Exp. | Exp. SD | log<sub>2</sub> of<br>Exp. | Exp. | Exp. SD | log<sub>2</sub> of<br>Exp. |
| **Fiber**<br>RS12070 | 0 | 1.00 | 0.15 | 0.00 | 1.01 | 0.18 | 0.01 |
|  | 1 | 1.30 | 0.31 | 0.38 | 3.85 | 0.62 | 1.94 |
|  | 2 | 1.45 | 0.34 | 0.53 | 277.05 | 58.02 | 8.11 |
|  | 4 | 1.87 | 0.48 | 0.90 | 62.66 | 17.56 | 5.97 |
| **Sheath**<br>RS12110 | 0 | 1.00 | 0.07 | 0.00 | 0.54 | 0.09 | -0.89 |
|  | 1 | 1.74 | 0.18 | 0.80 | 7.90 | 2.47 | 2.98 |
|  | 2 | 1.27 | 0.22 | 0.35 | 374.81 | 70.33 | 8.55 |
|  | 4 | 0.51 | 0.11 | -0.96 | 91.84 | 24.95 | 6.52 |
| **Tube**<br>RS12115 | 0 | 1.00 | 0.12 | 0.00 | 0.99 | 0.26 | -0.01 |
|  | 1 | 1.07 | 0.21 | 0.10 | 8.78 | 1.25 | 3.13 |
|  | 2 | 1.39 | 0.23 | 0.48 | 910.87 | 162.64 | 9.83 |
|  | 4 | 0.76 | 0.16 | -0.40 | 166.36 | 162.04 | 7.38 |

Exp. – normalized relative expression; Each value is averaged from 3 biological replicates; reference genes: *lpxC*, *rplU*
<sup>A</sup> Locus tag in the genome of *D. dadantii* 3937 (NC\_014500.1); shared prefix: DDA393

